# Identity domains in complex behavior: Toward a biology of personality

**DOI:** 10.1101/395111

**Authors:** Oren Forkosh, Stoyo Karamihalev, Simone Roeh, Mareen Engel, Uri Alon, Sergey Anpilov, Markus Nussbaumer, Cornelia Flachskamm, Paul Kaplick, Yair Shemesh, Alon Chen

## Abstract

Personality traits offer considerable insight into the biological basis of individual differences. However, existing approaches toward understanding personality across species rely on subjective criteria and limited sets of behavioral readouts, resulting in noisy and often inconsistent outcomes. Here, we introduce a mathematical framework for studying individual differences along dimensions with maximum consistency and discriminative power. We validate this framework in mice, using data from a system for high-throughput longitudinal monitoring of group-housed mice that yields a variety of readouts from all across an individual’s behavioral repertoire. We describe a set of stable traits that capture variability in behavior and gene expression in the brain, allowing for better informed mechanistic investigations into the biology of individual differences.

## Introduction

Individual differences are a hallmark of living organisms and central to our understanding of normal behavior and psychopathology. In humans, consistencies in emotional and behavioral expression have been extensively investigated and categorized by psychologists within the framework of personality traits^1,2^. In other species, however, the understanding of individual differences and the biological processes that underlie them has been hindered by the absence of a strong conceptual foundation behind the trait creation process and the lack of comprehensive behavioral screening paradigms.

Here we propose to resolve these issues using a computational framework for capturing and describing the space of individual behavioral expression by reducing diverse longitudinal behavioral data to trait-like dimensions. Personality traits can be thought of as having two crucial characteristics: (1) they capture and represent a continuous gradient of differences between individuals of the same species and (2) they tend to be stable for individuals over time. Thus, a mathematical formulation of a trait informed directly by these properties would be a dimension that captures the maximum behavioral variability between individuals while maintaining minimum variability within individuals over time. We use the term Identity Domains (IDs) to describe such traits obtained from decomposing a high-dimensional space of the measured behaviors. Conceptually similar to principal component analysis, which identifies the directions of maximum variability, our linear discriminant analysis (LDA) decomposition-based approach seeks the dimensions with maximal discriminative power and stability by maximizing the between- to within-individual variability ratio (Figure 1a). We validate this framework in mice, one of the most commonly used model organisms in neuroscience and psychiatry research, and a species that readily allows for exploration of the biological underpinnings of individual differences.

**Figure 1.**
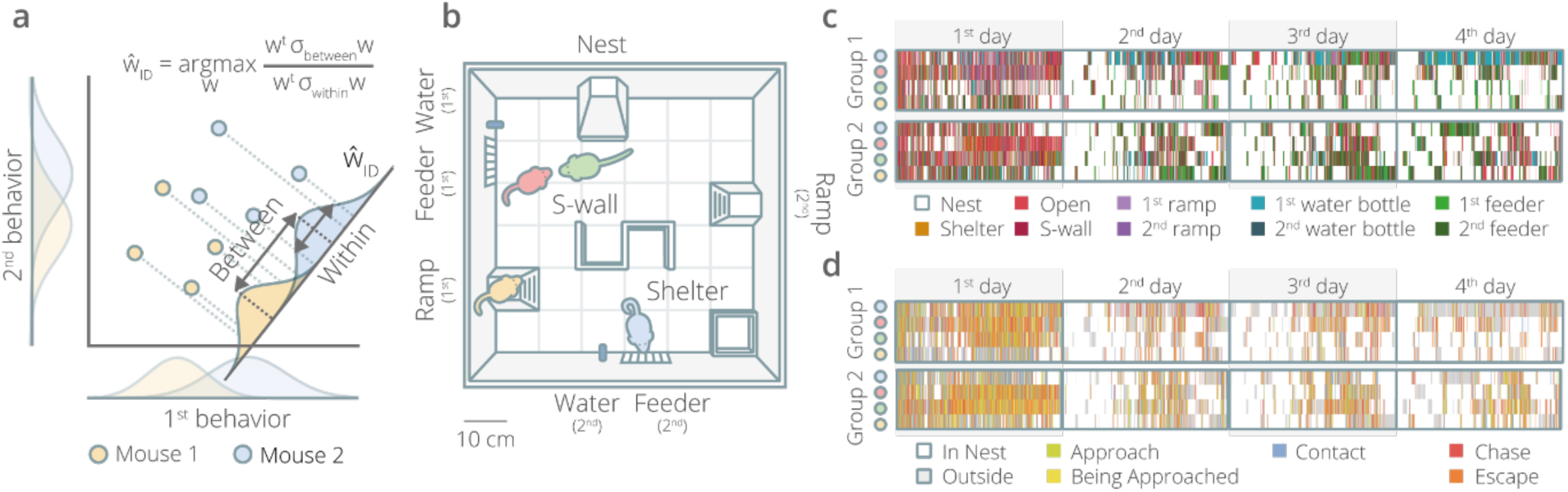
From behavior to personality. (a) The task of finding stable and discriminative trait-like dimensions can be formulated as an optimization problem. We used Linear discriminant analysis (LDA) to reduce the multidimensional behavioral space by creating dimensions that maximize the ratio of inter- to intra-subject variability. (b) Groups of four male mice, marked for tracking purposes with dyes of 4 different colors, were housed in an enriched environment where they could move and interact freely over multiple days. All of their movements were automatically tracked. Each arena contained a closed nest, 2 feeders, 2 water bottles, 2 ramps, an open shelter, and a S-shaped separation wall in the center. (c) Movement ethograms and (d) social ethograms for 2 representative groups of mice show intra-individual consistencies and inter-individual variability.

## Results

### The Social Box paradigm

In order to assess the broadest variety of ethologically relevant voluntary mouse behaviors, we used a long-term “Social Box” living paradigm, wherein mice are housed in an enriched, seminaturalistic environment in groups of four and monitored over multiple days ^3,4^ (Figure 1b-d, Supplementary Movie 1). Automatic location tracking of individuals allowed high-throughput behavioral data collection with readouts consisting of both individual (e.g., locomotion, exploration, foraging patterns) and social (e.g., approaches, contacts, chases) behaviors. A total of 60 features per mouse, per 12-hour active phase was collected (Supplementary Figure 1a). We initially monitored 42 groups of four outbred male mice (a total of 168 animals) left undisturbed over a period of at least four days.

### Linear Discriminant Analysis

An initial analysis of the readouts from this dataset revealed a subset of behaviors that, in themselves, were discriminating between and/or stable within individuals (Supplementary Figure 1b), suggesting that the Social Box paradigm could capture some of the information necessary for building IDs. We thus proceeded to train our algorithm on this dataset. Our analysis yielded four significant IDs that passed the threshold of less than 5% average overlap between individuals (Supplementary Figure 2, ID5 – the first dimension below this threshold is shown for comparison). The dimensions produced this way were uncorrelated, though not necessarily orthogonal, resulting in four IDs each spanning a very different behavioral subspace (Figure 2a).

**Figure 2.**
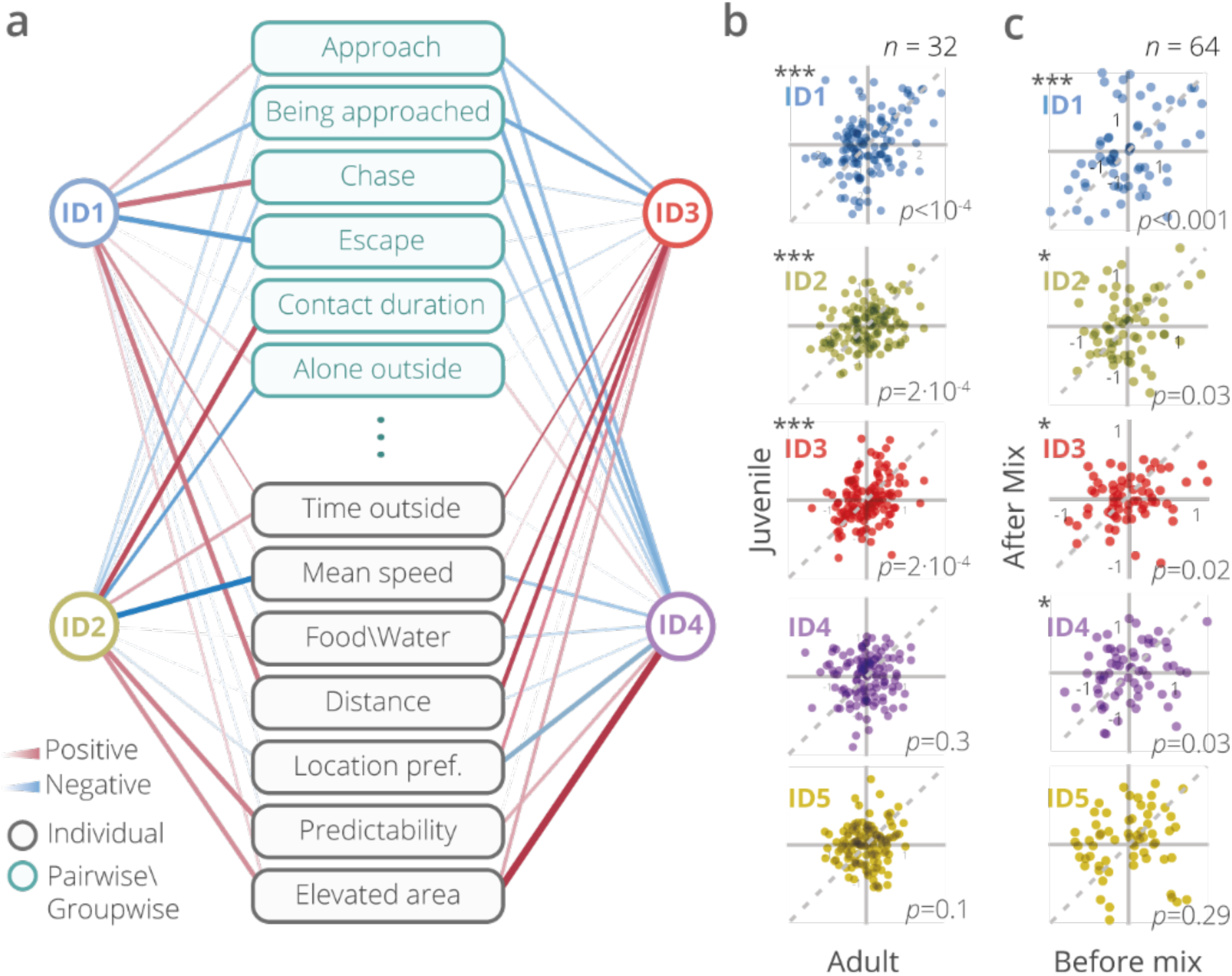
Testing the Identity Domains (IDs). (a) Running linear discriminant analysis (LDA) on the 60-dimensional behavioral space resulted in four significant IDs. The IDs are uncorrelated between themselves and are represented by multiple overlapping behaviors (showing 13 representative behaviors out of the 60, *see* Supplementary Figure 9). The width of connecting lines reflects the strength of the correlation (red – positive, blue – negative). (b) ID scores of mice as juveniles (4-5 weeks old) remained significantly stable at adulthood (15-16 weeks old) for the first three IDs. (c) ID scores of mice before and after being mixed into new social groups. Mice were significantly self-similar in their scores on IDs 1 through 4 across changing social environments. This relationship did not hold for ID5.

To test the replicability of the four IDs, we used a separate dataset composed of control mouse measurements (*n* = 208) in Social Boxes with a different layout (Supplementary Figure 3a), which yielded only a subset of the current behavioral readouts (37 different readouts per mouse per active phase). The scores on the top four IDs obtained from this dataset correlated strongly with the respective original scores (Supplementary Figure 3b). The strength of this relationship decreased steeply at ID5. We were thus able to replicate the initial ID structure on an independent dataset despite the differences in setup and readouts.

### IDs are stable over time, developmental stages, and social context

Having established our model, we proceeded to experimentally validate the IDs. To assess the stability of ID scores over time, we first tested their self-similarity from an average of the first four days of the experimental period to the 5^th^ day (Supplementary Figure 4). All IDs fulfilled this criterion. We then tested their stability over developmental time, by assessing juvenile mice (8 groups, 4-5 weeks old) in the social arenas. The same mice were tested again as adults (15-16 weeks old). Individual scores on IDs 1-3 were stable over this prolonged period of time (Figure 2b, Supplementary Figure 5), indicating that IDs assigned to juveniles captured individual differences that remained stable across developmental stages.

A major reason for the usefulness of behavioral traits over specific behaviors is that traits more closely approximate the intrinsic properties of an individual and are therefore more robust against manipulations of the social environment. Mice from 16 groups (64 individuals) that had been assigned ID scores based on the four-day baseline testing were then shuffled into new groups, such that no mouse had ever been exposed to any of its new group members and re-introduced to new arenas for another day of measurements (Supplementary Figure 6). For adult male mice, this is a dramatic and stressful manipulation causing significant changes in many of the behavioral readouts, especially those related to general locomotion and aggression. Despite these changes, the scores for IDs 1 through 4, but not ID5, remained significantly self-similar to their baseline state (Figure 2c). We additionally compared our model against PCA run on the same dataset. In this analysis only two out of the four top four principal components remain stable after this manipulation (Supplementary Figure 7a-c). Thus, mice tended to maintain the distinguishing individual characteristics captured by IDs despite substantial changes to the social environment.

### IDs combine information from a variety of standard behavioral tests

Having established that four IDs were stable over time and across social context, we set out to assess their ability to predict a range of standard behaviors typically measured in classical mouse behavioral paradigms. To this end, we submitted mice with known ID scores to a battery of established behavioral assays (Figure 3a-b). ID scores contained a significant portion of the information collected from classical tests (Figure 3a). The pattern of correlations between the various tests and ID scores additionally suggests that IDs represent complex entities that could not be fully captured without comprehensive behavioral screening. Moreover, these relationships contribute to the notion that IDs carry information about the hidden factors that are co-modulated across an animal’s behavioral repertoire. For example, ID1 was correlated with a measure of dominance in the social hierarchy (David’s score, Figure 5b) and also with features of locomotion in the open field test and memory recall in the object recognition test. All of these behavioral readouts appear to be expressions of a common underlying trait.

**Figure 3.**
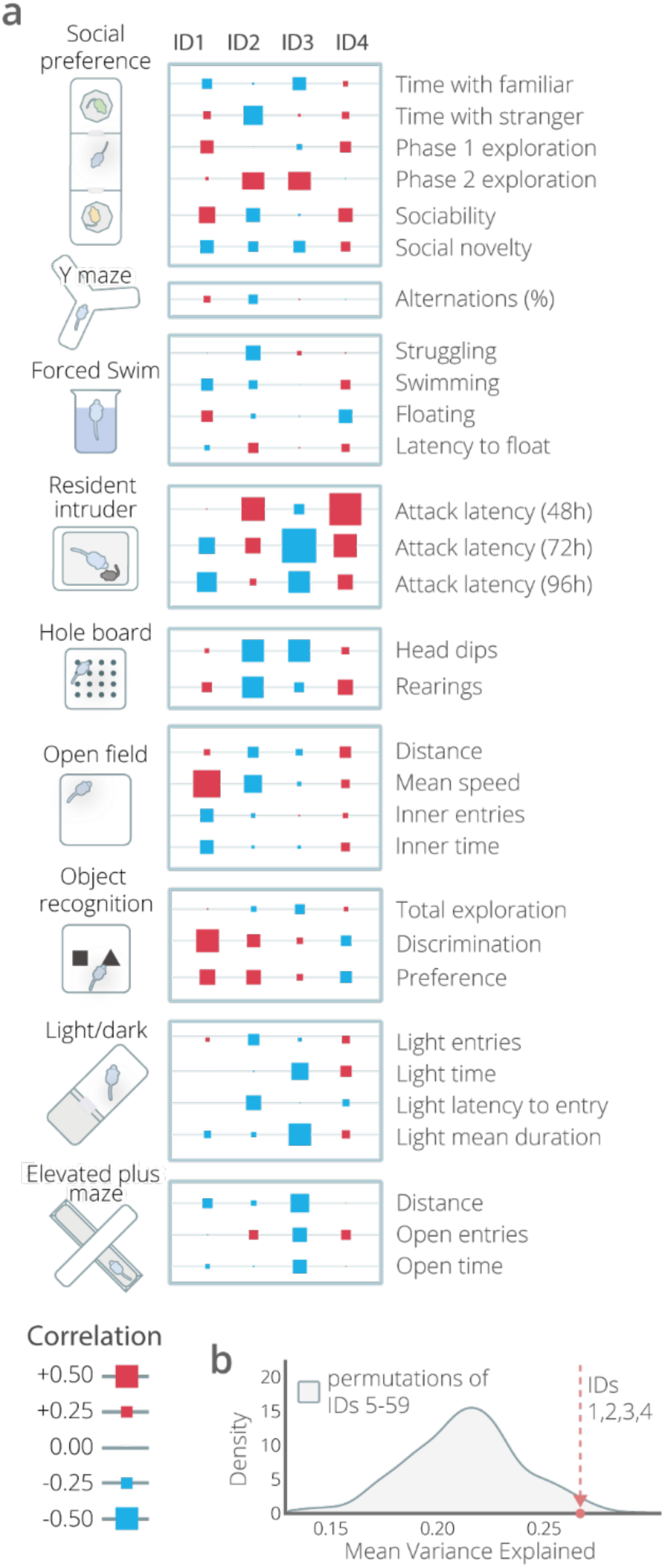
Identity Domains (IDs) are reflected in multiple standard behavioral tests. (a) ID scores predict multiple readouts across standard tests (two-tailed Pearson correlation statistic). Moreover, some standard test readouts are related to multiple IDs. The strength of each correlation is represented by the size of the squares (red – positive, blue – negative). (b) Variance explained by ID scores in standard behavioral assays. IDs 1-4 explain significantly more variance in “classical” behavioral test readouts than random sets of 4 from IDs 5-59.

**Figure 5.**
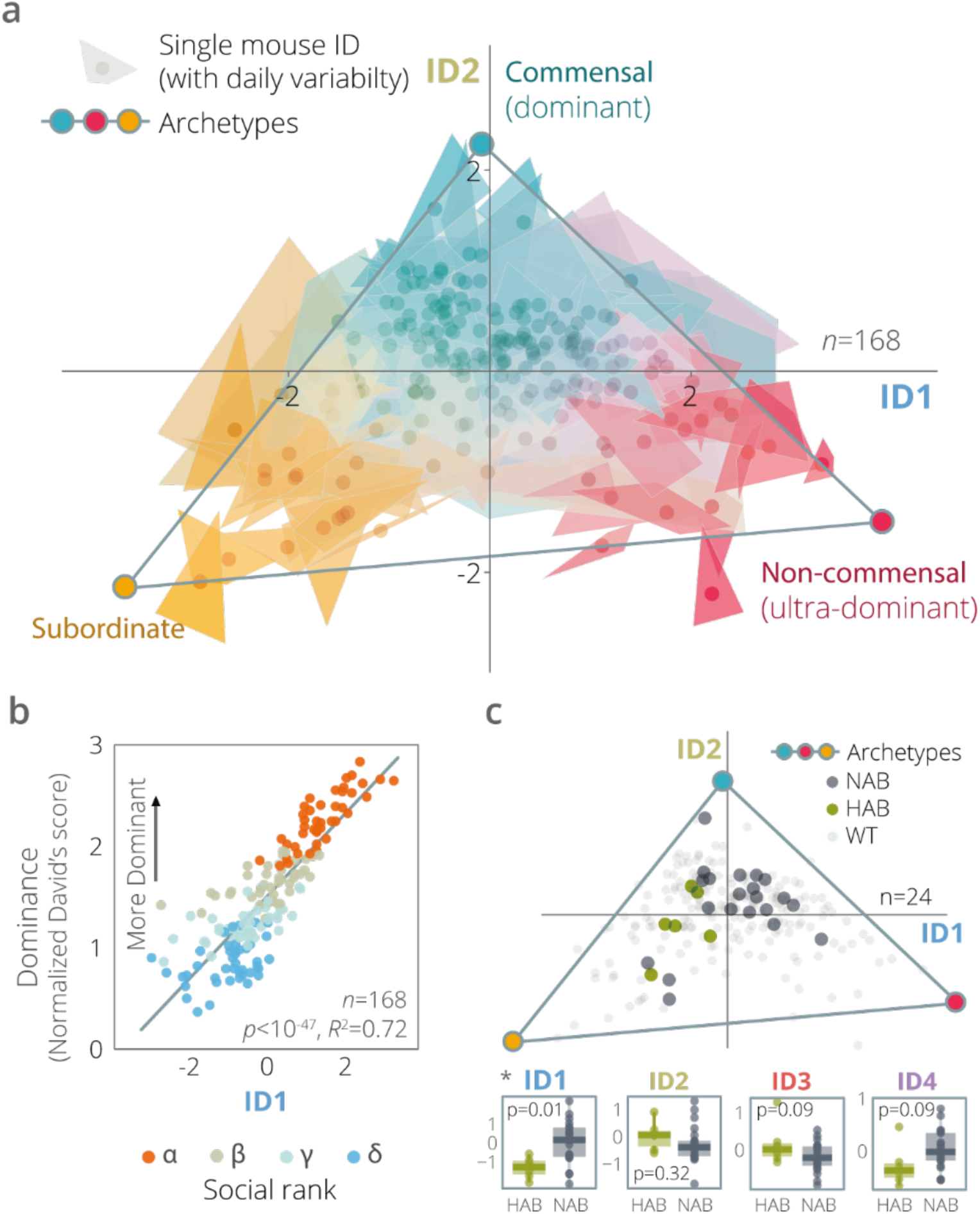
Personality space. (a) The personality space captured by the first two IDs forms a triangle with three archetypes at its corners. These archetypes may correspond to three behavioral strategies that mice exhibit in nature: commensal, non-commensal (ultra-dominant), and subordinate. The ID scores of each individual are represented by a trapezoid with scores on each day marking the four corners (a triangle is depicted if the fourth point falls inside the shape). Thus, the size of each trapezoid reflects the stability of ID scores for each individual over time (smaller means more stable). (b) ID1 scores predict social dominance levels (David’s Scores), calculated based on the number and directionality of aggressive interactions (*p* <10^-47^). (c) Mice selectively bred for high and normal anxiety-like behavior levels (HAB/NAB) mice were assessed using the IDs. A significant genotype effect was detected by ID1 and ID4. The Pareto space reveals that HAB mice tend to be more subordinate (upper panel).

### IDs capture transcriptomic variance in the brain

A major advantage of animal models is the ability to mechanistically investigate the link between brain and behavior. While the contributions of brain-transcriptomic differences to human personality traits remain largely unexplored due to major technical difficulties in performing such studies, mouse ID scores may prove a useful proxy. To assess whether ID scores captured transcriptomic variance in the brain, we performed bulk RNA-sequencing in mice that had been profiled in the Social Box (n = 32). For each individual, we sequenced three brain regions (Figure 4): the basolateral amygdala (BLA), the insular cortex (INS), and the medial prefrontal cortex (PFC), yielding a total list of 13,073 genes jointly detectable in all three regions. For each region, we assessed the average variance explained across the gene set by all four IDs and compared it against a distribution derived from shuffling the ID scores across individuals. Strikingly, in all three regions, the IDs performed significantly better in their true configuration than would be expected by chance, suggesting that ID score assignment is close to optimal with regard to their association with gene expression.

**Figure 4.**
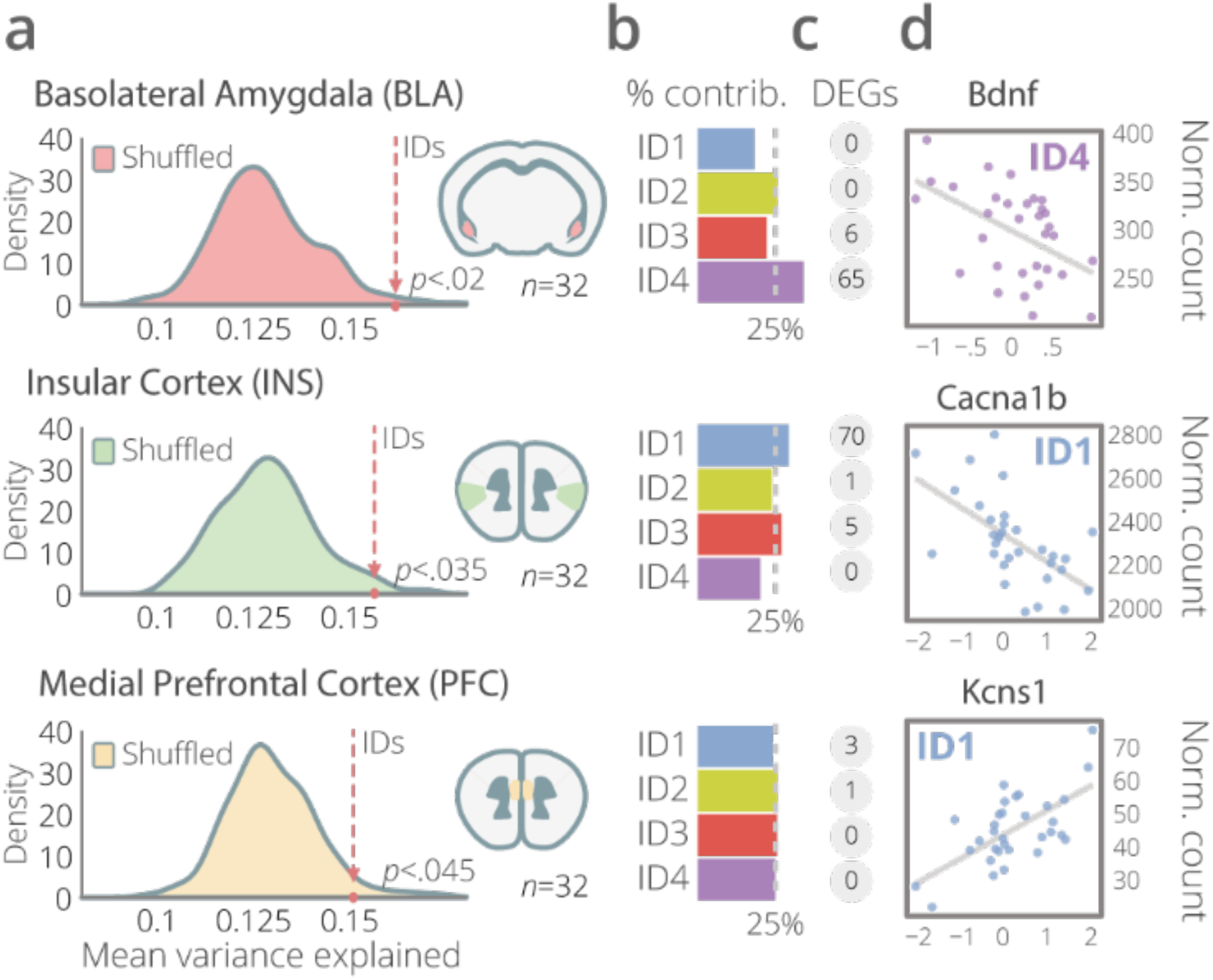
Identity Domains (IDs) carry information on gene expression in the brain. (a) RNA-sequencing results from the basolateral amygdala, insular, and medial prefrontal cortices of mice collected after baseline behavioral assessment. Plotted are the distributions of mean R-squared values for 200 models with four shuffled IDs as predictors. ID scores explain significant amounts of variance in the transcriptome (permutation test). (b) The relative contributions of each ID to the transcriptomic variance explained in each region. (c) Numbers of genes showing differential expression per ID after multiple testing correction. (d) Expression of one representative gene from each region plotted against individual scores on the ID with which it was associated.

### IDs discriminate between genotypes

Next, we tested the ability of IDs to capture and discriminate between individuals with known genetically driven differences in behavioral tendencies. For this purpose, we used the high-versus normal-anxiety (HAB/NAB) model, wherein mice are selectively bred over multiple generations for different levels of anxiety-like behavior on the Elevated Plus Maze test^5^ (Supplementary Figure 8a). We monitored heterogeneous groups composed of one HAB and three NAB individuals each in the Social Box and assigned ID scores to them. We were able to show that ID scores have considerable power in discriminating between the genotypes (Figure 5c, Supplementary Figure 7b).

### Personality Space

An important benefit that comes with having a known space of individual expression is the ability to search that space for points of biological interest, which may represent behavioral specializations. Using Pareto Task Inference, we found that ID1 and ID2 span a behavioral continuum on a triangle bounded by three personality archetypes (Figure 5a). Such a configuration can be interpreted as a tradeoff between 3 distinct evolutionary specializations, as previously shown for features of animal morphology^6^ and *C. elegans* locomotion^7^. Analogous archetypes were found in the replication dataset.

## Discussion

Personality is a complex entity that reflects stable individual differences and, in so doing, maps the space of phenotypic variability. Here we show that IDs provide a bias-free surrogate measure of personality obtained directly from behavioral data. IDs show considerable stability over time, developmental stages, and across social contexts. They allow quantitative exploration of personality differences in organisms in which such analyses were previously inaccessible. By drawing on consistent inter-individual differences, IDs capture the essence of personality, thus offering access to a biologically meaningful and evolutionarily relevant meta-behavioral

## Methods

### Animals

All animal experiments were approved by the Animal Care and Use Committees of either the Government of Upper Bavaria (Munich, Germany) or the Weizmann Institute of Science (Rehovot, Israel).

Male CD-1 (ICR) mice aged 8 to 12 weeks during the assessment were used for all experiments with the exception of the high-vs. normal anxiety-like behavior animals (HAB/NABs, see below). The animals were housed in an SPF facility in temperature-controlled rooms under standard conditions with a 12h light/dark cycle (lights on at 8 am). After weaning, the animals were housed in groups of four non-siblings per cage. At around 7-8 weeks of age, the mice were transferred to the behavioral testing rooms and painted. All animals were housed in temperature-controlled environment with food and water available *ad libidum*.

### Painting

The fur of each mouse was painted to enable identification by automatic video tracking. Painting was carried out under mild isoflurane anesthesia using commercially available semi-permanent hair dyes of three colors: Pillarbox Red, Voodoo Blue, and Sunshine (Tish & Snooky’s NYC Inc., New York). A fourth, green hue, was achieved by mixing the latter two dyes. The dyes were applied using a paint brush. Excess color was removed from the animal’s fur with tissues. The period under anesthesia was typically no longer than 10 min.

Mice were single-housed for several hours after painting and subsequently reunited with their cage mates. A minimum of 3 days of recovery/habituation was allowed following this procedure before the mice could be introduced into the social arenas.

### Social box setup

Mice were studied in a specialized arena designed for automated tracking of individual and group behavior. Each arena housed a group of four male mice. The arena consisted of an open 60 × 60 cm box and included the following objects: covered nest, open shelter, S-shaped wall, two water bottles, two feeders, and two elevated ramps. Food and water were available ad libitum. During the dark phase (12 hours) arenas were illuminated at 2 lux and during the light phase (12 hours) at 200 lux using LED lights. A color sensitive camera (Manta G-235C from Allied-Vision) was placed 1 m above the arena and recorded the mice during the dark phase. Mouse trajectories were automatically tracked offline using specially written software in Matlab (Mathworks Inc.).

In order to validate the identity domains (IDs) and ensure repeatability, we also computed the ID scores for mice which were recorded in arenas of a different design^3^. These alternative arenas were 75 × 50 cm and included a covered nest, closed shelter (which is smaller than the nest, and has only one entry), two elevated ramps, two feeders, a single water bottle, an elevated block that is away from the walls, and a Z-wall.

### Identification and classification of interactions between mice

We automatically identified and classified interactions between mice as events in which the distance between two mice (d) was less than 10 cm. We then used the movement direction of one mouse relative to another mouse (*θ*) to identify the nature of the contact for either of the mice. If for mouse A, the projection of the direction of its movement relative to mouse B was small enough 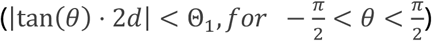 then it was considered as moving towards B; if 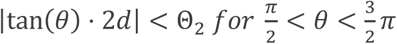 it was moving away from it; otherwise it was assumed idle with respect to the other mouse (Θ_1_ and Θ_2_ were found by optimization).

To classify aggressive and non-aggressive contacts, we first used a hidden Markov model^8^ to identify post-contact behaviors in which mouse A was moving towards B, and B was moving away from A (A was following B). We then used 500 manually labeled events to learn statistical classifiers of aggressive and non-aggressive post-contact behavior. For each event, we estimated a range of parameters, including individual and relative speed, distance, etc. and optimized a quadratic discriminant classifier^9^, a k-nearest neighbor algorithm based on these parameters, and a decision-tree classifier, that used these parameters at each tree intersection^10^. We found that for a test set of 1000 events, none of these classifiers were accurate enough individually, but that a combined approach in which we labeled an event as ‘aggressive’ if any of the classifiers labeled it as such – gave ∼80% detection with 0.5% false alarms.

### David’s score for dominance

We used the Normalized David’s score in order to assign each individual with a continuous measure of its social rank^11^. David’s score assumes a linear hierarchy where each pair from the group includes a more and a less dominant individual. The score is based on the measure of the fraction of interactions in which mouse *i* chased mouse *j* relative to the total number of agonistic interactions, which we denote as *P*_*ij*_. David’s score of each individual is the sum

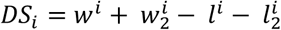

where *w*^*i*^ = Σ _*j* ≠ *i*_ *P*_*ij*_ is the sum of the fraction of times that mouse *i* has “won” (i.e., was the chaser), and 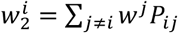 is a similar sum weighted by the *w*^*j*^ of the other mice, while *l*^*i*^ = Σ _*j* ≠ *i*_ *P*_*ij*_ is the sum of the fractions of “losses” (escapes), and accordingly 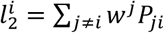 is its weighted sum. The score is then normalized to be between 0 and N-1 (where N is the number of subjects, which equals 4 in our case) by using the following formula:

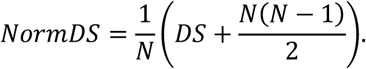

### Linear discriminant analysis

Linear discriminant analysis (LDA) is a method for finding a linear separator between two classes, or in its more general definition finding a subspace that best separates between multiple classes^9,12,13^. This subspace is obtained by finding a projection *W* which minimizes the Fisher-Rao discriminant defined as

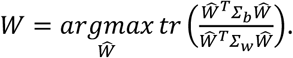

This solution to this optimization problem can be found by reformulating it using a Lagrange multiplier λ as

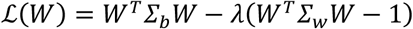

and solving

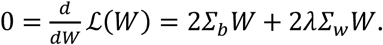

The solution to this equation is obtained by finding the top eigenvectors of 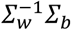.

### Fisher-Rao discriminant

The Fisher-Rao discriminant is used to measure how distinct two or more classes of samples are. The separation between classes is defined as the ratio of the variability between the classes Σ_b_ to the variability within the classes Σ_w_

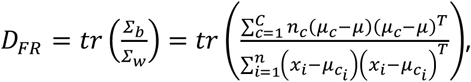

where *μ* is the global mean, 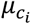 is the mean of the class associated with the *i*’th sample *x*_*i*_, *n*_*c*_ is the number of samples in class *c, n*, = *Σ*_*c*_ *n*_*c*_ is the total number of data samples, *C* is the number of classes, and tr(…) is the trace function. The larger this ratio, the better the discriminability of the classes. Note that the sum of the within- and between-variability is proportional to the total covariance of the data (*Σ*)

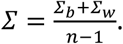

### Identity domains

Personality cannot be measured directly but it can be inferred from the behavior. We used the measured behavior of the mice for each of the four days they spent in the Social Box. The normalized measured behaviors of mouse *m* on day *d* are denoted by a vector *x*_*m,d*_ of dimension 60 (the number of behaviors; explanation of the normalization procedure follows). The ID *I*_*m*_ defines a distribution on the behavior space in the following way

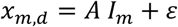

where *A* is a matrix linking the IDs to the behaviors, and *ε* is a distribution term (or noise due to variability or external factors). In order to estimate the IDs, we need to reverse this equation and find a *W* that would give us

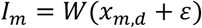

Note that since *A* is not usually square *W* is not simply a reversal of *A*.

In order to find *W*, which in turn would give us the IDs, we used LDA. Here the variability-within is defined as the variability of the same individual mouse on different days, or

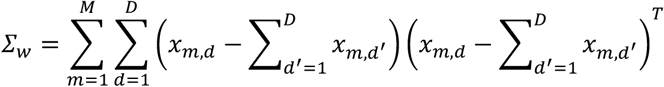

where *M* is the total number of mice (*M*=168), and *D* is the total number of days (*D*=4). Accordingly, the variability-within is the variability between mice or

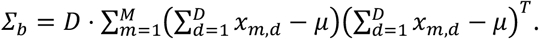

The link between the behavior and the IDs is obtained in the same way as in the classical LDA by solving

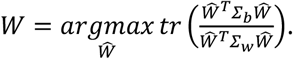

Once we have W, we can also find A by solving it as a linear regression problem of the form

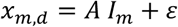

In order to avoid batch effects and drifts of the data we used quantile normalization on each behavior for each batch on each day separately. We quantile-normalized the data to have a normal distribution by computing the quantile of each sample and then computing the inverse of the normal cumulative distribution function (also known as the ‘probit’ function). If two or more samples were identical prior to the normalization they were all assigned the same value after normalization.

### Pareto optimality

In order to survive and reproduce, animals are constantly confronted with tasks such as finding food or evading predators. Often there is a tradeoff between tasks, so that the success of an animal in one task has to come at the expense of its performance on another. Recent work has shown that the best phenotypes are the weighted average of archetypes, which are phenotypes that specialize in one task^6,14^. These phenotypes can either be morphological, as for the beak sizes of ground finches, or behavioral phenotypes.

The shape of the phenotype space is determined by the number of archetypes, or the number of tasks the animal faces: In case of two archetypes, optimal phenotypes would fall on the line connecting the two archetypes, while if there are three archetypes, the phenotypes would be contained inside a triangle, and so on.

One direct outcome of this theory is that looking at the whole phenotype space makes it possible to deduce the location of the archetypes and thereby the different biological challenges that an animal faces. The positions of the archetypes are found using a hyperspectral un-mixing algorithm. The data is first centered to have zero mean, and then projected using principal component analysis into a subspace with dimension n – 1 (where n is the number of archetypes). Then an un-mixing algorithm is used in order to fit an n vertices polytope that best fits the data. Here we used minimal volume simplex analysis (MVSA), which is suitable for relatively small datasets since it does not allow for outliers. The analysis was performed in Matlab using the Pareto Task Inference (ParTI) package.

### Social box behavioral readouts

We collected a total of 60 different readouts for each mouse on each day. Due to the linearity of LDA, some of the behaviors we measured were computed with several different normalizations. The most common normalizations we used were: the total time in the arena (12 hours; abbreviated total), the time outside the nest (outside), and for interactions the total number of contacts (contacts).

#### Pairwise

- Time outside [1]: Fraction of time that the mouse spends outside of the nest. Normalizations: Total time (%).
- Frequency of visits outside [2]: The rate at which the mouse exits the nest. Normalizations: Total time (1/hour).
- Foraging correlation [3]: The correlation between the times that the mouse is outside the nest and the times that another mouse is outside the nest, averaged over all mice. For example, the foraging correlation between two mice would equal one if the mouse is always outside the nest when the other mouse is outside, and also enters the nest whenever the other mouse enters the nest. The correlation would be -1 whenever the mouse is outside, the other mouse is inside the nest. Normalizations: [3] none (au).
- Contact rate [4, 5]: Number of contacts the mouse had. A contact is defined as two mice being less than 10 cm apart while both are outside the nest. Normalizations: [4] Total time (1/hour), [5] Time outside (1/hour).
- Time in contact [6]: Fraction of time that a mouse is in contact with other mice while outside the nest. Normalizations: [6] Time outside (1/hour).
- Median\Mean contact duration [7, 8]: Median or mean duration of contacts. The contact duration does not include the times when the mouse approached, moved away from, or chased the other mouse. Normalizations: [7, 8] none (sec)
- Follow [12, 18, 24]: A follow is a contact that ended with one mouse going after another mouse until disengagement. Follows can be either aggressive (chases) or non-aggressive. Normalizations: [12] Number of contacts (au), [18] Time outside (1/hour), [24] Total time (1/hour).
- Being-followed [13, 19, 25]: Number of times a mouse is followed at the end of a contact. It can be either in an aggressive (chases) or non-aggressive manner. Normalizations: [13] Number of contacts (au), [19] Time outside (1/hour), [25] Total time (1/hour).
- Chase [10, 16, 22]: Chases are interactions that ended with the mouse pursuing another mouse in an aggressive manner. Aggressiveness was determined using a classifier that was trained on labeled samples (see methods). Normalizations: [10] Number of contacts (au), [16] Time outside (1/hour), [22] Total time (1/hour).
- Escape [11, 17, 23]: Number of time that the mouse was aggressively chased by another mouse. Normalizations: [11] Number of contacts (au), [17] Time outside (1/hour), [23] Total time (1/hour).
- Non-aggressive follow [14, 20, 26]: Number of times the mouse has followed another mouse at the end of a contact in a non-aggressive way. Normalizations: [14] Number of contacts (au), [20] Time outside (1/hour), [26] Total time (1/hour).
- Non-aggressively being-followed [15, 21, 27]: Number of times the mouse was followed by another mouse at the end of a contact in a non-aggressive way. Normalizations: [15] Number of contacts (au), [21] Time outside (1/hour), [27] Total time (1/hour)
- Approach [28, 29, 30, 31, 32]: An approach is a directed movement of the mouse towards another mouse that ends in contact. Not all interactions necessarily start with an approach, while others might start mutually with both mice approaching each other. Normalizations: [28] none (au), [29] Time outside (1/hour), [31] Number of contacts (au), [32] total (1/hour), [30] Time outside with one or more mice (1/hour).
- Being-approached [33, 34, 35]: Number of times the mouse was approached by another mouse. Normalizations: [33] Number of contacts (au), [34] Time outside (1/hour), [35] none (au).
- Approach-escape [36]: Fraction of contacts in which the mouse initiated the contact and ended up being chased. Normalizations: [36] Number of aggressive contacts (au).
- Difference between approaches and chases [9]: The total number of chases is subtracted from the total number of approaches. Normalizations: [9] none (au).

#### Individual

- ROI exploration [37, 38]: Quantifies the amount of exploration the mouse is doing. Measured as the entropy of the probability of being in each of the 10 regions-of-interest (ROIs). Mice that spend the same amount of time in all regions will get the highest score, while mice that spend all their time in a single ROI will be scored zero. When normalized to the time outside, the computation of the entropy differed also by ignoring the probability of being inside the nest. Normalizations: [37] none (bits), [38] Time outside (bits/hour).
- Grid exploration [59]: Quantifies the amount of exploration the mouse is doing. Analogously to ‘ROI Exploration’, grid exploration was determined using entropy, however, instead of looking at the ROIs, we divided the arena into a 6x6 grid (10 cm by 10 cm; a total of 36 possible locations). Normalizations: [59] none (bits).
- Predictability [60]: Measures how predictable the paths that the mouse takes as the mutual information between its current and previous location in the arena. For that, the arena was divided into a 6x6 grid (10 cm by 10 cm; a total of 36 possible locations), and for each cell we computed the probabilities of it moving to any of the adjacent cells. Normalizations: [60] none (bits).
- Distance [58]: The total distance traveled by the mouse while outside the nest. To smooth the tracking, the mice locations were sampled once every second. Normalizations: [58] none (m).
- Median\Mean speed [54, 55]: Median or mean speed while outside the nest. To smooth the computation of the speed, the locations of mice were sampled once every second. Normalizations: [54, 55] none (m/sec).
- Tangential velocity [56]: The tangential component of the speed, or the part of speed perpendicular to the previous direction of movement. Normalizations: [56] none (m/sec).
- Angular velocity [57]: The rate of change in the direction of the mouse. Normalizations: [57] none (rad/sec).
- Food or water [39, 40]: Time spent next to the feeders or water bottles. Normalizations: [39] Total time (au), [40] Time outside (au)
- Food [41]: Time spent next to the feeders. Normalizations: [41] Time outside (au).
- Water [42]: Time spent next to the water bottles. Normalizations: [42] Time outside (au).
- Feeder preference [43]: Time spent in the feeder adjacent to the nest (feeder 1) relative to the further-away feeder (feeder 2). Normalizations: [43] none (au).
- Water preference [44]: Time spent near the water bottle adjacent to the nest (water 1) relative to the further-away water bottle (water 2). Normalizations: [44] none (au).
- Elevated area [45, 46]: Time spent on an elevated object in the arena: ramps or block (in the other arena setting). Normalizations: [45] Total time (au), [46] Time outside (au).
- Open area [47]: Time spent in the open area (outside of the nest and any of the ROIs). Normalizations: [47] Time outside (au).
- Shelter [48]: Time spent in the shelter, which is a box closed only on its sides. Normalizations: [48] Time outside (au).
- Ramps [49]: Time spent on the elevated ramps. Normalizations: [49] outside (au).
- S-wall [50]: Time spent in the S-wall. Normalizations: [50] Time outside (au).
- Distance from walls [51]: Average distance from the walls while in the open area. Normalizations: [51] none (cm).
- Distance from nest [52]: Average distance from the nest (while outside of the nest). Normalizations: [52] none (cm).
- Alone outside [53]: Fraction of time the mouse is outside while all other mice are in the nest. Normalizations: [53] Total time (au).

### Standard behavioral assays

All behavioral tests were performed several days after the social arena assessment in the same test room in two batches of 8 groups each using two different test sequences. During this time, the mice were housed in their original groups.

Timeline 1 consisted of the open field (OF)/novel object recognition (NOR) test, social preference test (SPT), and dark-light transfer (DaLi) test. Timeline 2 consisted of the OF/NOR, followed by the elevated plus maze (EPM), DaLi, SPT, and forced swim (FST) tests. In both cases, each test was followed by a minimum of 48 hours of rest.

#### Open field and novel object recognition tests

The OF and NOR tests were performed in 60 × 60 cm boxes under minimal illumination (2-3 lux) in three sessions. Each animal was introduced to the arena for 5 min, then briefly removed and reintroduced for 5 more minutes to the same arena, now with two identical objects placed at predetermined locations. Finally, each animal was reintroduced to the arena after a 4 h delay for 5 minutes with one of the two identical objects replaced by a novel one^15^. Object preference was calculated as the novel/familiar object ratio, while the discrimination index was calculated as the ratio of preference to phase 2 total exploration time.

#### Dark-light transfer and elevated plus-maze tests

Anxiety-like behaviors were assessed using the DaLi or EPM tests performed in standard behavioral apparatuses. The illumination of the light sections of each apparatus was set at 200 lux and the duration of both tests was set to 5 min.

#### Forced swim test

Behavioral despair was measured using the FST. Each animal was placed in a 2 L transparent beaker filled halfway with room-temperature water for 6 min. Floating, swimming, and struggling times were manually scored by experienced observers.

#### Social preference test

The SPT was performed under low illumination (2 lux) over three sessions in a three-chamber apparatus. This consisted of a middle chamber connected to two chambers on each side by a door. An empty metal grid cone was placed in the center of each of the two side-chambers. During the first session, the doors to the side chambers were closed and each test mouse was introduced into the middle chamber and allowed to habituate to it for 5 min. The doors were subsequently opened and a stimulus CD-1 mouse was placed under one of the metal grid cones, the other remaining empty, for 10 minutes. Sociability was calculated based on this session as the ratio of time spent in the chamber with the stimulus mouse to time in the chamber with the empty cone, weighted by total time spent in both of these chambers. Finally, in a following 10 min session, a different stimulus mouse was placed under the other cone. Preference for social novelty was assessed using the ratio of time with novel versus familiar mouse weighted by total time in either chamber.

#### Y-maze alternation task

Working memory was assessed using the Y-maze alternation task. Each mouse was introduced for 5 min into a Y-shaped three-arm apparatus with distinguishing visual cues on the walls at the end of each arm. The proportion of spontaneous non-repeated subsequent entries into each of the three arms (alternations) from the total number of three-arm entries (including repeat entries) was used as the final readout.

#### Resident-intruder test

Aggression toward an unfamiliar intruder mouse was assessed using a resident-intruder paradigm. For this test, each mouse was single-housed in a fresh type 2 cage. At each time point, 48, 72, and 96 h after single-housing, an intruder C57/BL6 mouse was introduced into the cage. Latency to first aggressive interaction was assessed by an observer. Each trial was interrupted after the first overtly aggressive confrontation or after 15 min.

#### Holeboard exploration test

Exploratory behavior was measured in the holeboard exploration test. Each mouse was introduced for 5 min into a 40 × 40 cm arena surrounded by transparent Plexiglas walls with 4 × 4 equally spaced holes on the floor. Number of head dips into any hole as well as number of rearings during the test interval were assessed by an observer.

### RNA-sequencing

Brains were dissected from animals sacrificed with an overdose of isoflurane and flash-frozen. Tissue samples were cryopunched using a 1 mm diameter punching tool at Bregma 1.98 mm (medial prefrontal cortex (PFC), insular cortex (INS), 600 um depth) or an 0.8 mm diameter punch at Bregma -0.7 mm (basolateral amygdala (BLA), 1000 um depth) according to Paxinos and Franklin, 1998. Total RNA was isolated using the miRNeasy micro kit (QIAGEN) after homogenization by a Bullet Blender (Next Advance). Residual genomic DNA was removed using the Turbo DNA-free kit (Ambion®, Invitrogen, CA, USA). RNA integrity and absence of DNA was confirmed by an Agilent RNA ScreenTape (4200 TapeStation, Agilent, eRIN all >7.8) and Qubit DNA High sensitivity kit, respectively. Sequencing libraries were prepared using the Illumina TruSeq Stranded Total RNA Library Preparation HT Kit using mammalian RiboZero Gold following the standard protocol starting from 300 ng (PFC), 500 ng (INS), and 375 ng (BLA) of total RNA using 12 cycles of PCR amplification. Libraries were quality-checked using Bioanalyzer DNA High Sensitivity chips (Agilent Technologies, St. Louis, MO) and quantified using the KAPA Library Quantification Kit (KAPA Biosystems, Boston, MA). Sequencing was performed on 6 lanes of an Illumina HiSeq4000 PE 2x100 (Illumina, San Diego, CA) multiplexing all samples.

Sequencing was performed on a HiSeq4000 generating 100 base pair paired-end reads. Read quality was checked using FastQC^16^ and subsequently adapters were trimmed using cutadapt^17^. For quantification of transcript expression levels, Kallisto^18^ was executed using Gencode M11 annotation and collapsed to gene level.

Count data was pre-filtered for low counts at a threshold of ≥ 20 counts per sample in a minimum of 31 out of 32 samples per region. In addition, the top three most highly expressed genes were excluded, resulting in a total list of 13,073 genes. Differential gene expression analyses were performed in DESeq2^19^. Heteroscedasticity in the data was reduced using the DESeq2 regularized logarithm transformation. The plate row of each sample was identified as a potential batch effect and corrected for using the limma package^20^.

The normalized and log-transformed count data was then used as the outcome of a linear model with either the real or the shuffled ID scores (day 1 in the social box) as predictors (*rlog(count) = ID1 + ID2 + ID3 + ID4 + ε*). Total variance explained by the model, as well as the fraction of variance explained by each individual predictor were estimated using the variancePartition package^21^. Two hundred models with shuffled ID scores were run per region to generate a distribution of mean variance explained across all genes and the model with the real ID scores was tested against this distribution using a one-sample t-test.

### Code availability

All the code used in the LDA implementation in MATLAB will be made available upon request. Likewise, the self-similarity tests implemented in MATLAB, as well as the R code used in the RNA-seq data analysis will be made available upon request.

## Data availability

The datasets generated during and/or analyzed during the current study are available from the corresponding author on reasonable request.

## Supplementary Information

Supplementary Figures 1-9

Movie 1: A representative segment from a video recording of the social arenas with a group of four colored mice. Overlaid on top of the video are illustrations of tracked mouse movements and the layout and components of the social arena.

## Acknowledgments

We thank Noa Eren, Iain Couzin, Carsten Wotjak, and Chadi Touma for their assistance, advice, and constructive criticism. Thanks to Jessica Keverne for professional English editing, formatting, and scientific input. Our thanks also to Ori Maoz for his unique insights into the mathematics their interpretation. Finally, we would like to extend special thanks to the recently passed Chaya Tannor for fascinating discussions on human personality.

A.C. is the head of the Max Planck Society - Weizmann Institute of Science Laboratory for Experimental Neuropsychiatry and Behavioral Neurogenetics. This work is supported by: an FP7 Grant from the European Research Council (260463, A.C.); a research grant from the Israel Science Foundation (1565/15, A.C.); the ERANET Program, supported by the Chief Scientist Office of the Israeli Ministry of Health (A.C.); the project was funded by the Federal Ministry of Education and Research under the funding code 01KU1501A (A.C.); research support from Roberto and Renata Ruhman (A.C.); research support from Bruno and Simone Licht; I-CORE Program of the Planning and Budgeting Committee and The Israel Science Foundation (grant no. 1916/12 to A.C.); the Nella and Leon Benoziyo Center for Neurological Diseases (A.C.); the Henry Chanoch Krenter Institute for Biomedical Imaging and Genomics (A.C.); the Perlman Family Foundation, founded by Louis L. and Anita M. Perlman (A.C.); the Adelis Foundation (A.C.); the Marc Besen and the Pratt Foundation (A.C.); the Irving I. Moskowitz Foundation (A.C.). S.K. is supported by the International Max Planck Research School for Translational Psychiatry (IMPRS-TP).

## Author contributions

O.F and S.K. designed the experiments, analyzed the results, and wrote the manuscript. M.N., C.F., and P.K. assisted in experiments. M.E. performed the sample collection and RNA-sequencing library preparation. S.R. performed the preprocessing of the RNA-sequencing data and contributed to the final analyses. U.A., S.A., and Y.S. contributed to the manuscript. A.C. supervised and supported the project.

The authors declare no competing financial interests.

---

**Supplementary Figure 1.**
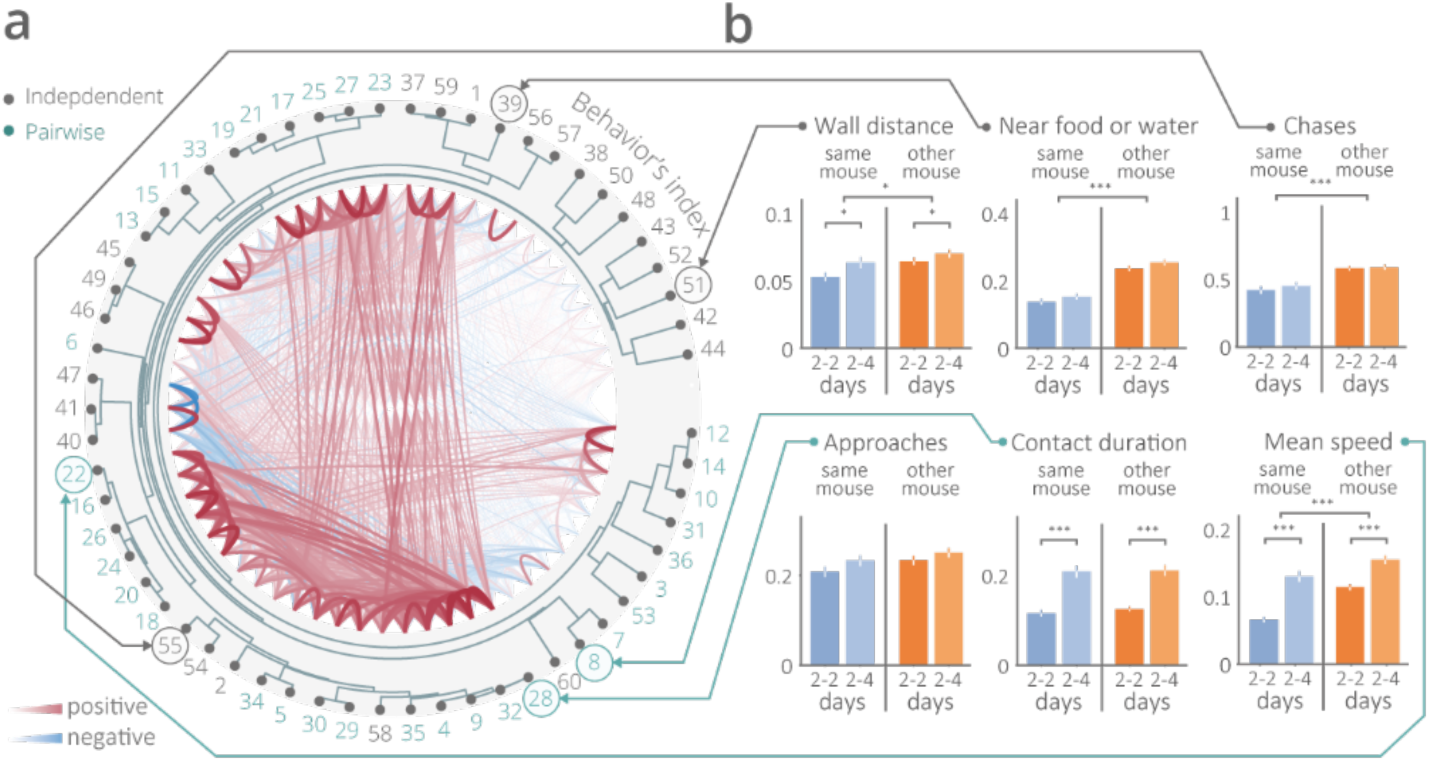
Individual differences and consistencies. (a) Behavioral readout structure. Hierarchical clustering and cross-correlations of the 60 behavioral readouts. Behavioral readouts tend to cluster based on whether they are independent (related to 1 mouse) or pairwise (derived from the locations of 2 mice). (b) Some behavioral parameters were consistent within individuals over time (e.g., approaches, chases), while some parameters could discriminate between individuals (e.g., mean speed, wall distance; \*\*\**p*<0.001, \**p*<0.05).

**Supplementary Figure 2.**
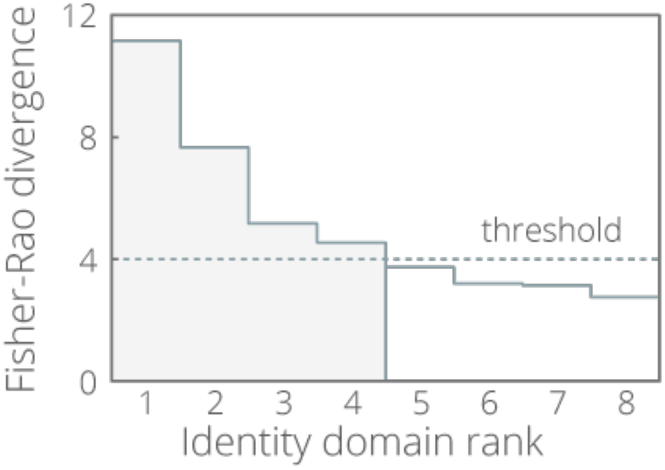
Between-within variability ratio. Identity domain (ID) components ranked by their Fisher-Rao coefficient. Four components had a Fisher-Rao-score below 4, indicating a greater contribution of between over within-individual variability.

**Supplementary Figure 3.**
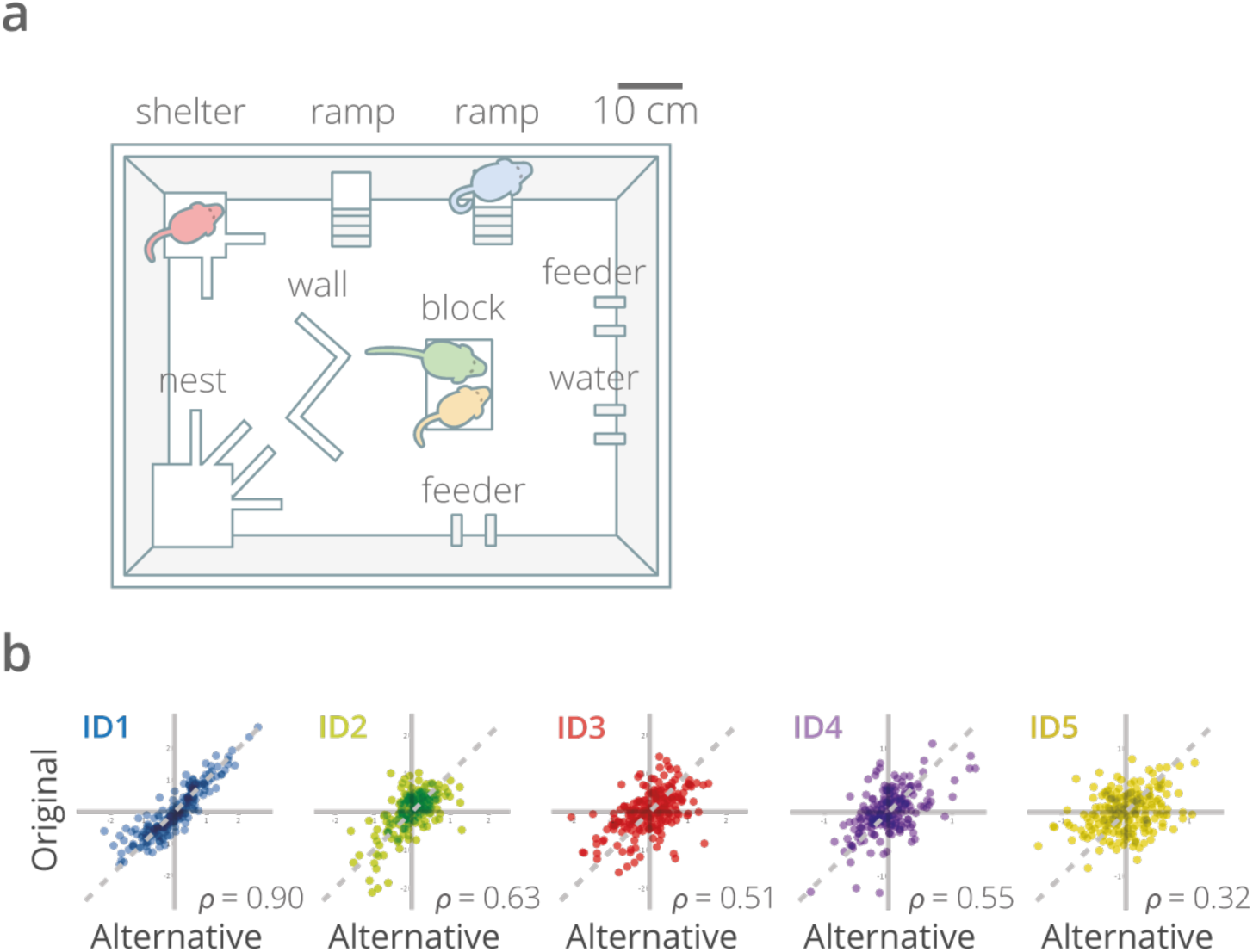
Validation of the identity domains (IDs) in a second dataset from an alternative setup. (a) Alternative social arena (50 × 70 cm) with a different locations and types of objects compared to arena shown in Figure 1 (b). (b) IDs 1-4 show intermediate to strong correlations between the original and replication datasets.

**Supplementary Figure 4.**
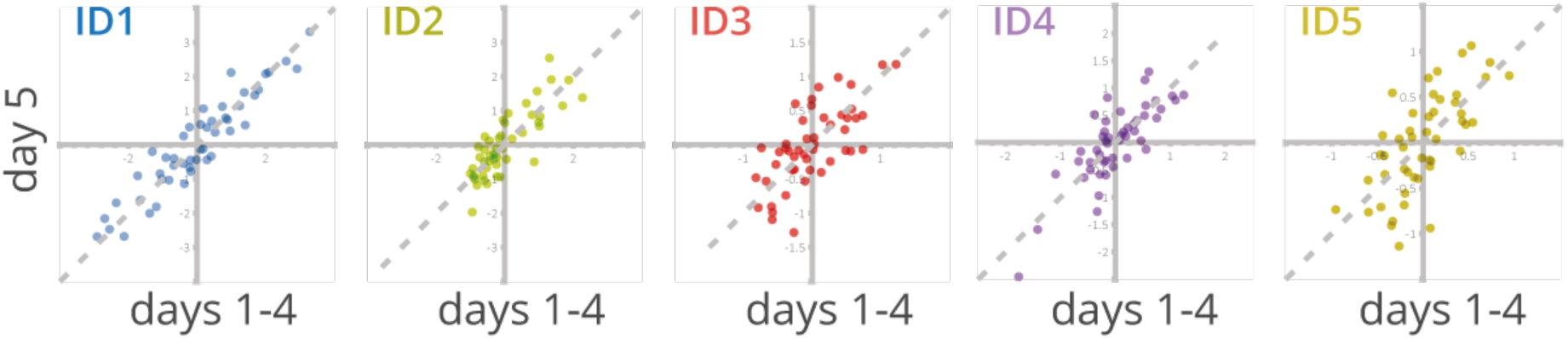
Identity domain (ID) stability over a short timescale. IDs were stable over experimental time, such that average ID scores for experimental days 1 through 4 could predict the corresponding scores for each animal on day 5.

**Supplementary Figure 5.**
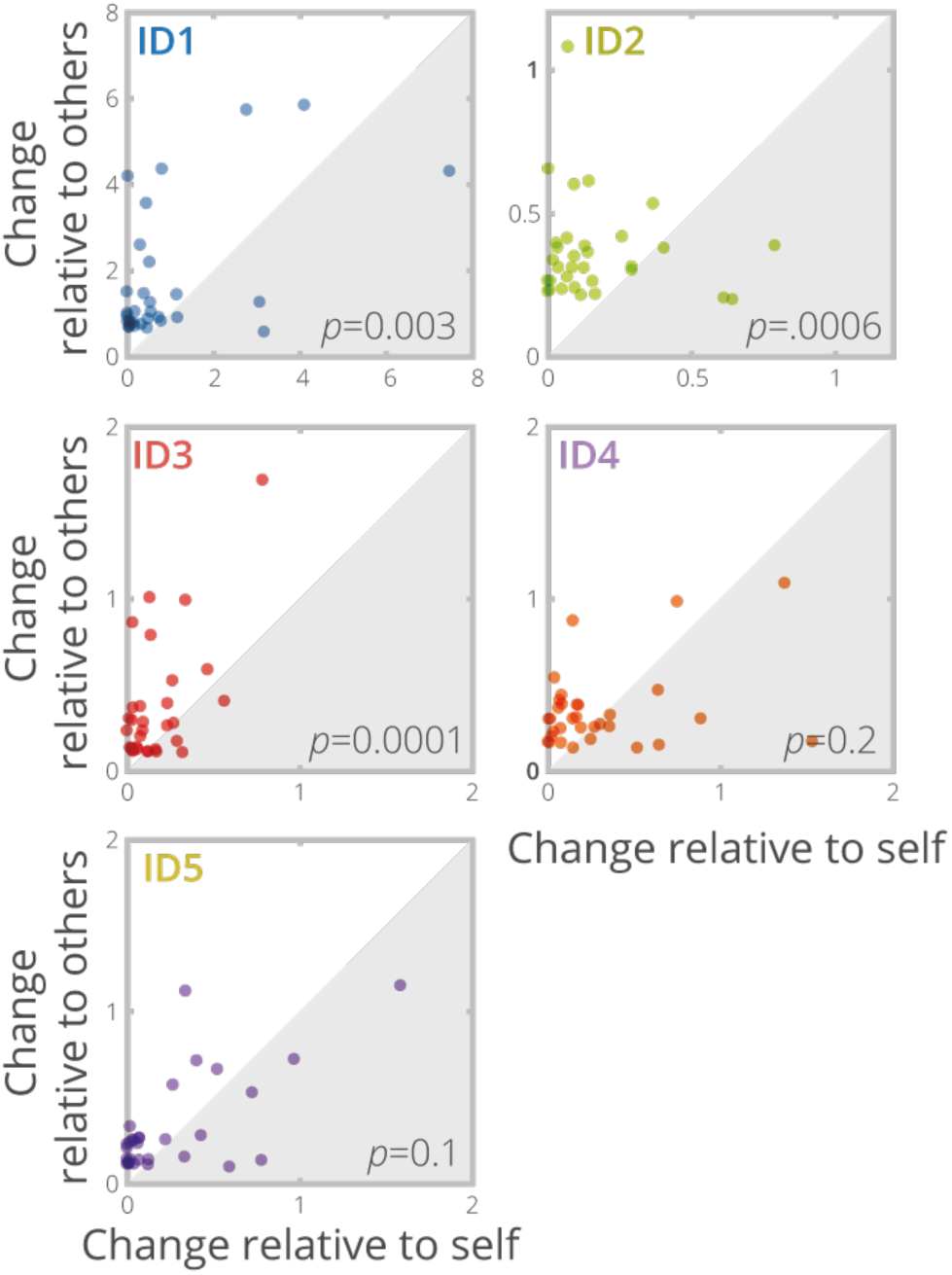
Change over time with respect to self or others. ID stability during aging was tested by comparing the ID scores of individuals measured once juveniles (4-5 weeks old) and once more during adulthood (15-16 weeks old). Depicted here are change in ID scores relative to one’s own initial score versus relative to the scores of all other individuals. Points in the shaded region represent greater individual changes, whereas points in the unshaded region represent changes that were larger relative to other individuals than to oneself.

**Supplementary Figure 6.**
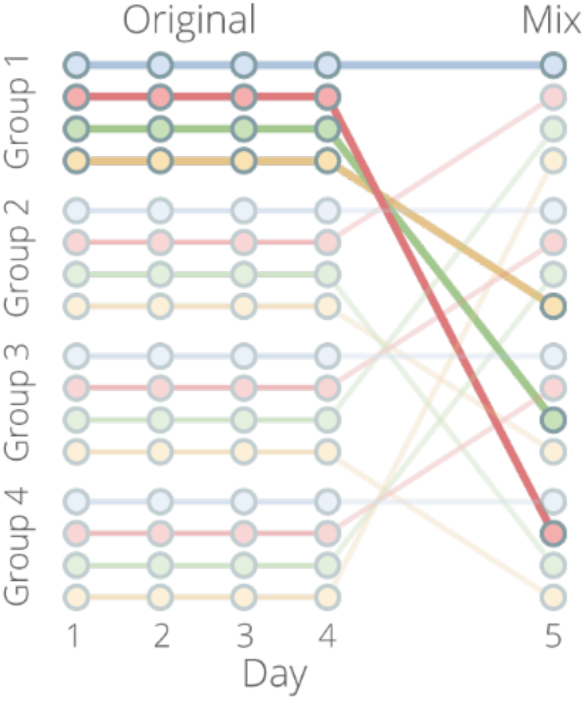
Group shuffle diagram. Mice were observed in the social boxes over 4 days and re-grouped on day 5 such that no mouse was familiar with any of its new conspecifics (*n* = 64, 16 groups).

**Supplementary Figure 7.**
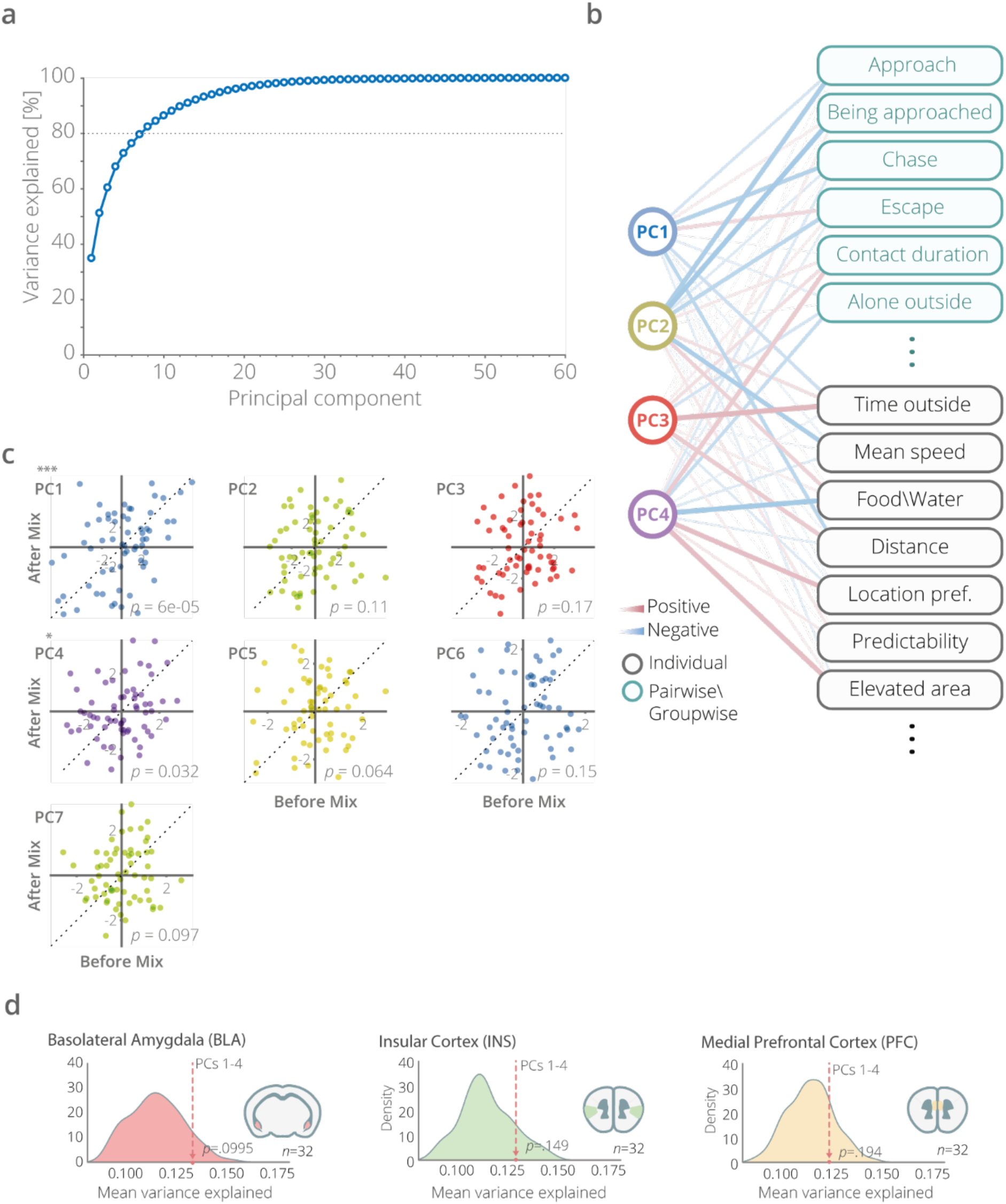
Principal components analysis (PCA) on the initial set of behaviors. In order to compare how LDA performs relative to better-known and more commonly used dimension reduction method, PCA was performed on the same initial dataset as used to generate the IDs. (a) The percent variance of the behavioral data explained by each principal component (PC). (b) Correlations between scores on each PC and an abbreviated list of behavioral readouts. (c) The stability of PC scores was tested as with the IDs before and after mixing the mouse groups such that all individuals were unfamiliar to one another. Only the first principal component remained stable after the mix. (d) Scores on the first four PCs were used as predictors of transcriptomic variance in RNA-sequencing data from three different brain regions. This analysis directly mimicked the equivalent analysis performed using the four IDs (PC scores from day 1, 200 shuffled PC score sets). The top four PCs did not carry more overall transcriptomic information than would be expected by chance.

**Supplementary Figure 8.**
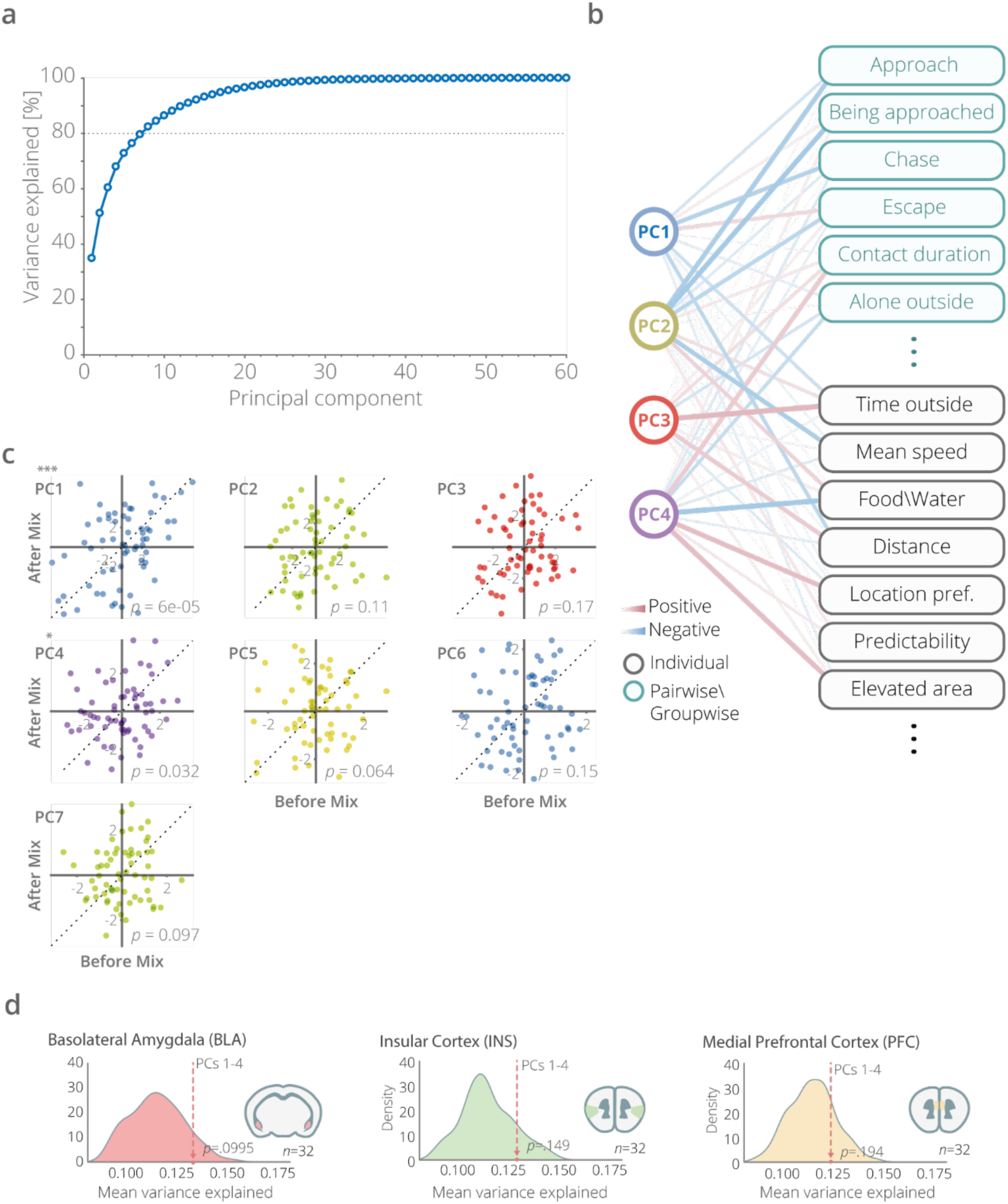
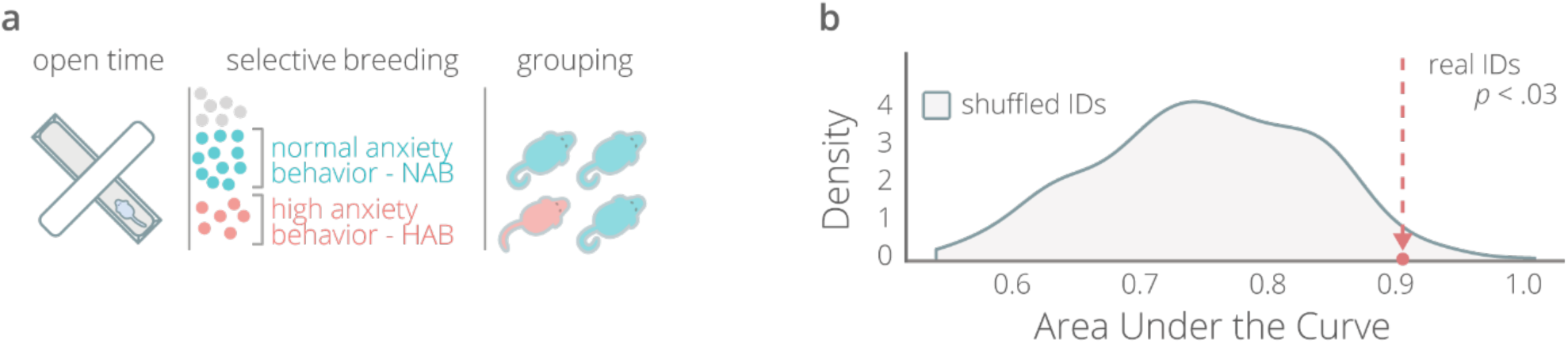
High-anxiety (HAB) versus normal-anxiety (NAB) mouse model. (a) Selective breeding for high versus normal anxiety-like behavior levels (HAB/NAB) was performed for over > 40 generations starting with outbred CD-1 mice^5^. Selection was based on results of the Elevated Plus Maze test (% time in the open arm). After the animals of each respective genotype were weaned, they were mixed into groups of three NABs and one HAB each. (b) The power of the identity domain (ID) scores to detect genotype was tested directly using the area under the receiver operating characteristic curve of a model predicting genotype based on IDs 1-4. The area under the curve of this model was compared against a distribution created based on 200 trials with shuffled ID scores.

**Supplementary Figure 9.**
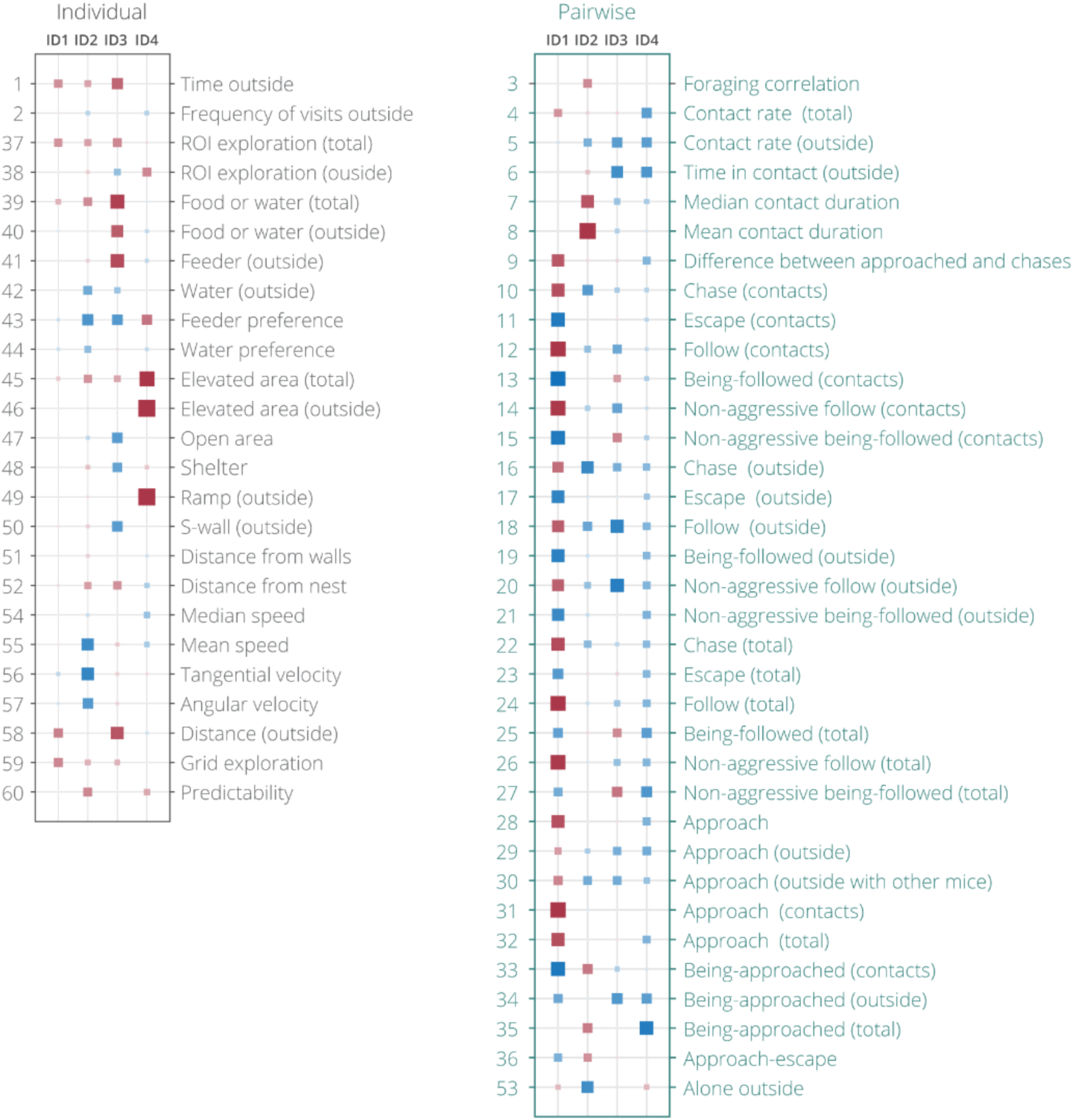
Correlations between identity domains (IDs) and their contributing behavioral readouts. The readouts are separated into individual (based on the movements of a single mouse) and pairwise (based on the movements of a mouse and one more of its group members).

